# Enhancers of human and rodent oligodendrocyte formation predominantly induce cholesterol precursor accumulation

**DOI:** 10.1101/2022.06.21.497032

**Authors:** Joel L Sax, Samantha N Hershman, Zita Hubler, Dharmaraja Allimuthu, Matthew S Elitt, Ilya Bederman, Drew J Adams

## Abstract

Regeneration of myelin in the CNS is being pursued as a potential therapeutic approach for multiple sclerosis. Several labs have reported small molecules that promote oligodendrocyte formation and remyelination *in vivo*. Recently, we reported that many such molecules function by inhibiting a narrow window of enzymes in the cholesterol biosynthesis pathway. Here we describe a new high-throughput screen of 1,836 bioactive molecules and a thorough re-analysis of more than 60 molecules previously-identified as promoting oligodendrocyte formation from human, rat, or mouse oligodendrocyte progenitor cells (OPCs). These studies highlight that an overwhelming fraction of validated screening hits, including several molecules being evaluated clinically for remyelination, inhibit cholesterol pathway enzymes like EBP. To rationalize these findings, we suggest a model that relies on the high druggability of sterol-metabolizing enzymes and the ability of cationic amphiphiles to mimic the transition state of EBP. These studies further establish cholesterol pathway inhibition as a dominant mechanism among screening hits that enhance human, rat, or mouse oligodendrocyte formation.

## 1. INTRODUCTION

Myelin is a lipid-rich substance found in the mammalian central nervous system (CNS) that is critical for saltatory conduction and structural support of neurons.^1–3^ Oligodendrocytes are the resident myelinating cells of the CNS and are lost in demyelinating diseases, most notably, multiple sclerosis (MS).^4, 5^ A CNS stem cell population known as oligodendrocyte precursor cells (OPCs) can differentiate into new oligodendrocytes that are able to replenish lost myelin in MS.^6–8^ Although OPC differentiation into oligodendrocytes occurs in many patients with demyelinating disease, this process is limited and ultimately fails to curb disease progression.^1^ Future therapeutics that function to drive OPC differentiation into oligodendrocytes to replace damaged myelin could improve patient prognosis.^9, 10^

To identify starting points for the discovery of future remyelinating therapeutics, various labs have used phenotypic screens of FDA-approved drugs and other bioactive small molecules to identify dozens of enhancers of oligodendrocyte formation.^11–16^ Several of these small molecules also enhance remyelination in animal models of MS, and one, clemastine, recently induced statistically significant improvement of visual evoked potential latency in optic neuritis patients.^17^ While validated hits from these screens span a diverse range of canonical biological targets, we have previously shown that a broad swath of such molecules modulate oligodendrocyte formation not by their canonical targets but by inhibition of a specific range of cholesterol biosynthesis pathway enzymes—CYP51, sterol 14-reductase, and emopamil-binding protein (EBP) (Figure S1).^15, 18^ Inhibition of these enzymes leads to accumulation of their 8,9-unsaturated sterol substrates, which are necessary and sufficient for enhanced oligodendrocyte formation.^15^

While many validated hits are established to induce 8,9-unsaturated sterol accumulation to drive oligodendrocyte formation, it remains unclear whether this finding is consistent across the broad swath of OPC differentiation assay conditions that have been used throughout the field. In particular, various labs have assayed mouse, rat, or most recently human OPC differentiation to oligodendrocytes; have tested OPCs that in some cases include disease-causing mutations; and have used screening libraries at various concentrations. In this study, we sought to evaluate the generality of 8,9-unsaturated sterol accumulation among oligodendrocyte-enhancing small molecules obtained across this diversity of screening contexts. To do so, we performed a new screen using low concentrations of bioactive small molecules and performed sterol profiling in mouse OPCs and human OPC-like glioblastoma cells on more than 60 validated hits identified by multiple labs using a range of OPC sources, genetic backgrounds, and screening methods. Notably, the first set of hits obtained via screening of human OPCs also predominantly induced 8,9-unsaturated sterol accumulation in human OPC-like glioblastoma cells, indicating that these cholesterol precursors also appear to be a leading mechanism by which existing small molecules promote human oligodendrocyte formation. Our studies have revealed that a large majority of small-molecule enhancers of oligodendrocyte formation identified by this set of screens function by inhibiting EBP in OPCs, with smaller numbers of validated hits instead targeting other cholesterol pathway enzymes or the thyroid hormone receptor, another well-validated remyelination target. Additional analyses establish multiple rationales for why this range of cholesterol biosynthesis enzymes are so frequently targeted by small-molecule screening hits, providing important context for this dominant mechanism for promoting oligodendrocyte formation and remyelination.

## 2. RESULTS AND DISCUSSION

### 2.1 Screening at lower concentrations highlights a new EBP inhibitor and thyroid hormone analogs as enhancing oligodendrocyte formation

Previously we used our established high-content imaging assay measuring the differentiation of mouse OPCs to mature MBP+ oligodendrocytes to screen 3,000 bioactive small molecules at 2 µM to identify enhancers of oligodendrocyte formation. Analysis of the top 10 novel hits from this screen revealed that all ten inhibited the cholesterol biosynthesis enzymes CYP51, sterol 14-reductase, or EBP.^15^ In this study, we re-evaluated a subset of 1,836 molecules within our bioactives collection (Selleck Bioactive Compound Library-I; L1700) at a substantially lower screening concentration of 100 nM. The goal of screening at this lower concentration was to identify potent enhancers of oligodendrocyte formation that may have been overlooked in past screens, for example due to cell death at micromolar concentrations. In brief, we treated OPCs in 384-well plates for 72 h with each bioactive small molecule at 100 nM and subsequently immunostained for myelin basic protein (MBP) to identify mature oligodendrocytes (see Methods). At this screening concentration, six small molecules strongly enhanced oligodendrocyte formation (Figure 1A). Four of these top enhancers of oligodendrocyte formation were analogs of thyroid hormone, which is well-established to drive oligodendrocyte formation via the thyroid hormone receptor.^19^ A fifth hit was amorolfine, which we previously established functions by inhibition of sterol 14-reductase in OPCs.^15^ Our screen also revealed one potent enhancer of oligodendrocyte formation, clomifene, a selective estrogen receptor modulator (SERM) (Figure 1A-1C).^20^ While clomifene enhanced MBP^+^ oligodendrocyte formation in the low nanomolar range, cytotoxicity and a reduction in oligodendrocyte formation was observed at concentrations at or above 1 µM (Figure 1C). Clomifene’s cytotoxicity in the micromolar range likely obscured its ability to enhance oligodendrocyte formation in previous high-throughput screens, further validating our 100 nM screening approach. Multiple SERMs have previously been shown to inhibit EBP.^15^ We next used GC-MS based sterol profiling (see Methods) to characterize clomifene’s effects on sterol synthesis in mouse OPCs at the same concentrations as assayed for OPC differentiation (Figure 1C and 1D). We observed accumulation of EBP’s substrate sterols zymostenol and zymosterol at similar concentrations to clomifene’s maximal effects on OPC differentiation, indicating a phenotypic correlation between clomifene’s ability to inhibit EBP and drive oligodendrocyte formation (Figure 1D and 1E). Overall, this screen again highlighted EBP and thyroid hormone signaling as common modes of action among high-throughput screening hits for enhancing oligodendrocyte formation.

**Figure 1.**
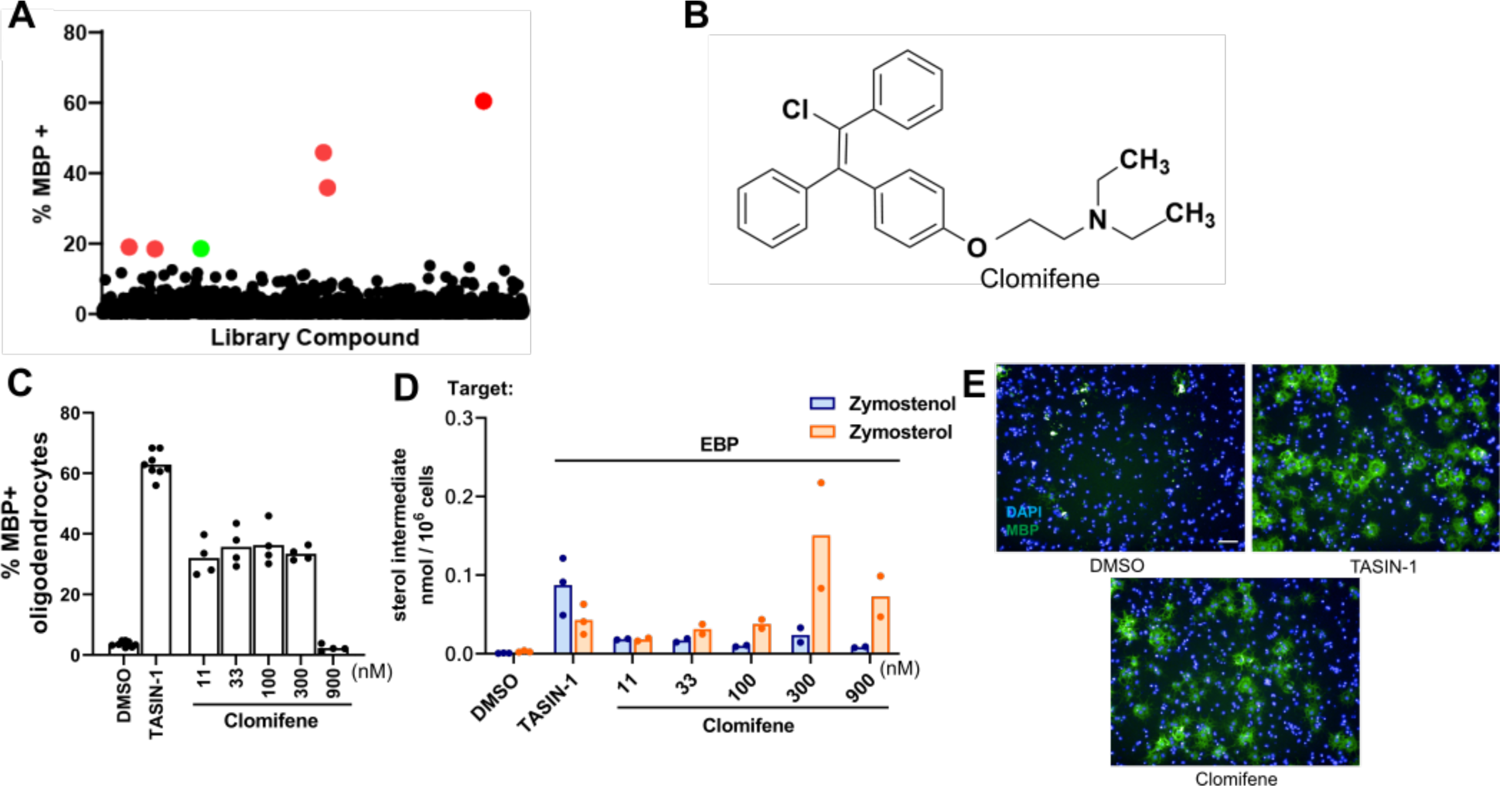
High-throughput screen reveals novel and known EBP inhibitors and thyroid hormone analogs as potent enhancers of oligodendrocyte formation. (A) Percentage of MBP+ oligodendrocytes generated from mouse OPCs following treatment with a collection of over 1,800 bioactive small molecules at a uniform screening dose of 100 nM for 72 h. Red dots represent previously characterized enhancers of oligodendrocyte formation. The green dot represents clomifene. (B) Chemical structure of clomifene. (C) Percentage of MBP+ oligodendrocytes generated from mouse OPCs following treatment with doses of clomifene for 72 h. *n* ≥ 4 wells per condition. (D) GC-MS based quantification of zymostenol and zymosterol levels in mouse OPCs treated with clomifene at the concentrations shown for 24 h. *n*=2 wells per condition. For (C), (D), DMSO and 1 µM TASIN-1, a known EBP inhibitor, served as the negative and positive control, respectively. (E) Representative images of mouse OPCs treated with DMSO, clomifene at 300 nM, and TASIN-1 at 1 µM. Nuclei are labeled with DAPI (blue), and oligodendrocytes are shown by immunostaining for MBP (green). Scale bar, 100 µm. (C) and (D) represent two independent experiments.

### 2.2 Past and newly-validated hits identified using stem cell-derived mouse OPCs inhibit cholesterol biosynthesis to induce 8,9-unsaturated sterol accumulation

In our previous screen of over 3,000 bioactive small molecules, we found that each of the top ten novel enhancers of oligodendrocyte formation were inhibitors of CYP51, sterol 14-reductase, or EBP and induced accumulation of 8,9-unsaturated sterols at the screening dose.^15^ In this study, we further evaluated 16 compounds that fell just below our previous hit threshold. We initially tested each molecule at the previous HTS concentration of 2 µM and found 6 of the 16 molecules effectively promoted oligodendrocyte differentiation, defined as at least a 1.5-fold increase in myelin basic protein-positive mature oligodendrocytes (Figure 2A, 2G and S2A). To capture molecules that might be effective at slightly higher concentrations, we retested initially ineffective molecules at 6 µM, which yielded 4 additional enhancers of oligodendrocyte formation (Figure 2B and S2B).

**Figure 2.**
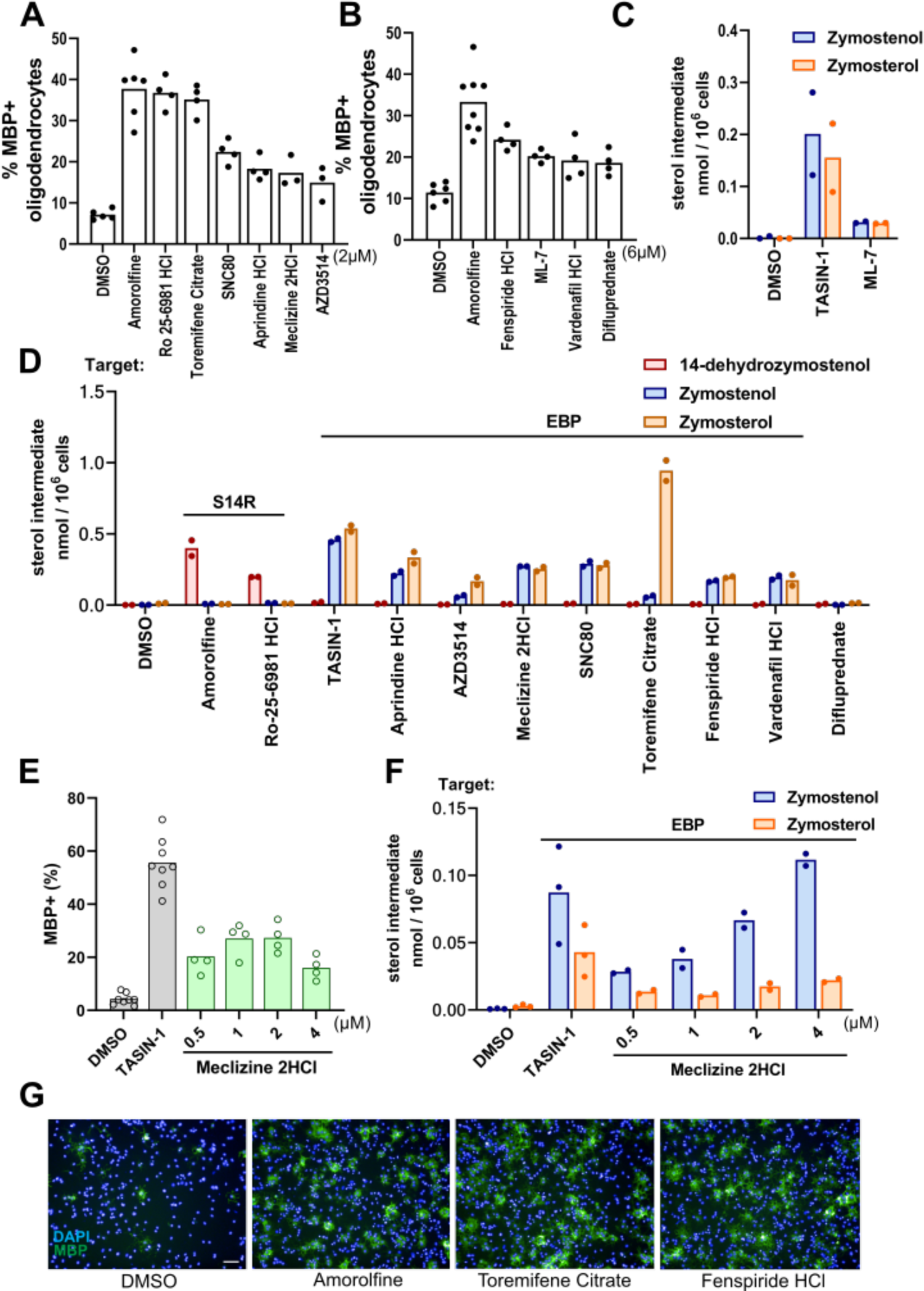
EBP and sterol 14-reductase are common enzyme targets in screening hits. (A), (B) Percentage of MBP+ oligodendrocytes generated from mouse OPCs following treatment with bioactive small molecules at 2 µM (A) and 6 µM (B). *n*=4 wells per condition, except DMSO and amorolfine, a known sterol 14-reductase inhibitor, *n*=8 wells, with >1,000 cells analyzed per well. (C), (D) GC-MS based quantification of 14-dehydrozymostenol, zymostenol, and zymosterol levels in mouse OPCs treated for 24 h with TASIN-1 (1 µM), a known EBP inhibitor, and the indicated screening hits at the concentrations shown in panel (A) and (B). *n*=2 wells per condition. (E) Percentage of MBP+ oligodendrocytes in mouse OPCs treated with meclizine at the indicated concentrations. (F) GC-MS based quantification of zymostenol and zymosterol levels in mouse OPCs treated with meclizine in dose response at the indicated concentrations. *n*=2 wells per condition. (G) Representative images of mouse OPCs treated with amorolfine at 600 nM, toremifene citrate at 2 µM, and fenspiride at 6 µM. Nuclei are labeled with DAPI (blue), and oligodendrocytes are shown by immunostaining for MBP (green). Scale bar, 100 µm. For percentage of MBP+ oligodendrocytes for all bioactive small molecules that were screened, see Fig. S2. S14R, sterol 14-reductase.

For these 10 small molecules that enhanced oligodendrocyte formation, we used GC-MS-based sterol profiling to measure inhibition of cholesterol pathway enzymes by quantitating accumulation of the sterol substrates of these enzymes (Figure 2C-D). When tested at the effective screening dose for each molecule (2 or 6 µM), nine of the 10 hits led to sterol accumulation with 8 inhibiting EBP and 1 inhibiting sterol 14-reductase. Only difluprednate, a glucocorticoid receptor agonist, showed no evidence of cholesterol pathway inhibition. Glucocorticoid receptor modulators including clobetasol have previously been suggested to promote OPC differentiation via glucocorticoid receptor signaling, consistent with our finding of no effect on cholesterol biosynthesis.^16^ As expected, three molecules that did not enhance OPC differentiation showed no ability to induce 8,9-unsaturated sterol accumulation using GC-MS based sterol profiling (Figure S2A-C). Finally, we also tested one molecule, meclizine, in dose-response in both our OPC differentiation and GC-MS profiling assays and found general alignment between 8,9-unsaturated sterol accumulation and MBP+ oligodendrocyte formation (Figure 2E-F). Ultimately, our reanalysis of this small-molecule screen revealed 9 novel inhibitors of EBP or sterol 14-reductase that enhance oligodendrocyte formation, further supporting the centrality of these enzyme targets among small molecules identified in high-throughput screens.

An earlier screen of FDA-approved drugs in murine pluripotent stem-cell-derived OPCs identified miconazole, clobetasol, and many other drugs as enhancers of oligodendrocyte formation.^16^ Reanalysis of the 15 hit molecules that were validated as dose-responsive in this study revealed that 10 have subsequently been characterized as inhibiting CYP51, sterol 14-reductase, or EBP in OPCs, with an eleventh molecule (haloperidol) shown to bind to and inhibit mouse EBP in vitro (Figure 3A).^15, 21, 22^ We next evaluated whether the remaining four validated hit molecules might also inhibit these enzymes. Using GC-MS-based sterol profiling, we noted that three of the four hits induced accumulation of 8,9-unsaturated sterols, with three EBP inhibitors and one CYP51 inhibitor identified (Figure 3B-C). Clobetasol, another glucocorticoid agonist, was the only compound that did not inhibit cholesterol synthesis. Notably, this classifies 14 of the 15 previously validated hits in this screen as either EBP, CYP51, or sterol 14-reductase inhibitors.

**Figure 3.**
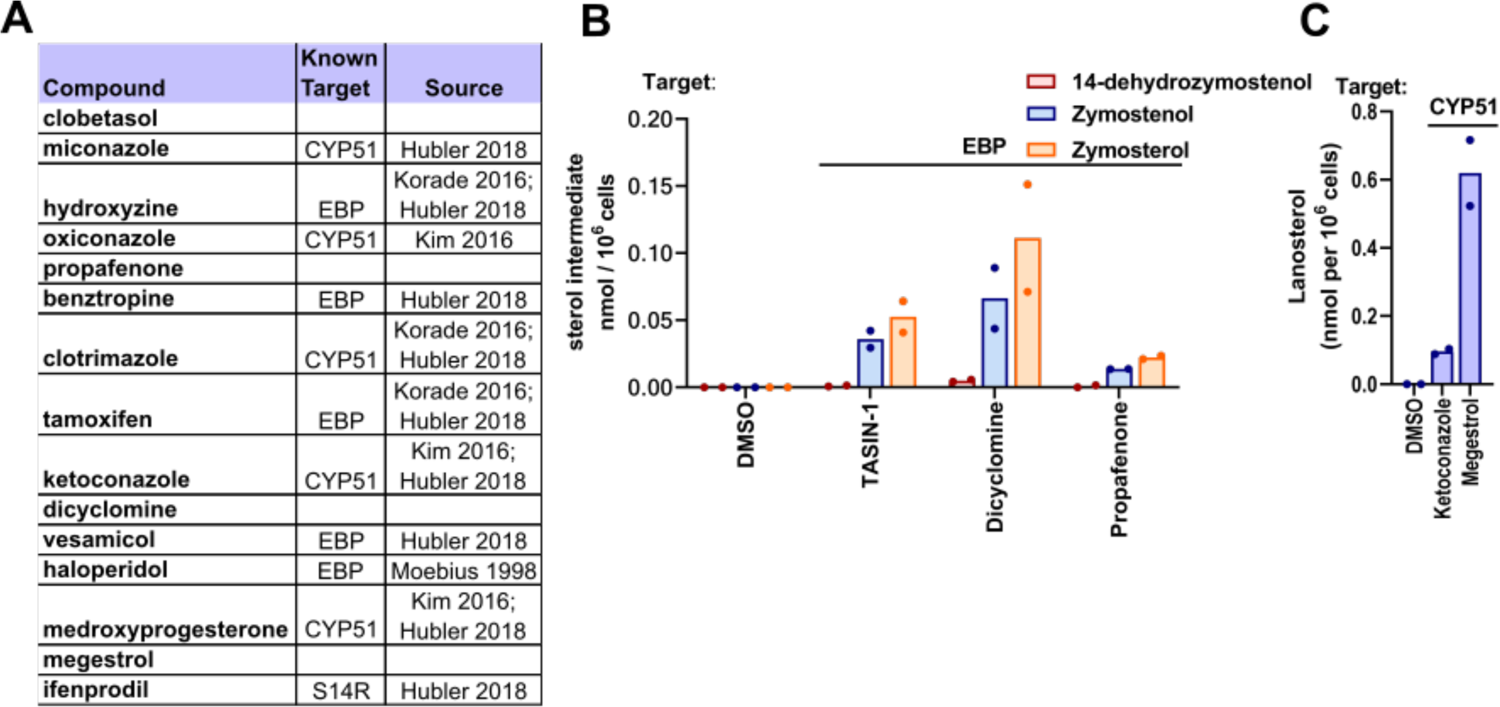
Remyelinating agents inhibit CYP51, EBP, and sterol 14-reductase. (A) Validated screening hits from Najm et al, 2015, with annotations of subsequently-established inhibitory activity for EBP, CYP51, or sterol 14-reductase. (B) GC-MS based quantification of 14-dehydrozymostenol, zymostenol, and zymosterol levels in mouse OPCs treated for 24 h with dicyclomine and propafenone at 2 µM. (C) GC-MS based quantification of lanosterol levels in mouse OPCs treated with ketoconazole (2.5 µM), a known CYP51 inhibitor, and megestrol (13.4 µM) for 24 h. n=2 wells per condition. S14R, sterol 14-reductase.

### 2.3 Top enhancers of *jimpy* oligodendrocyte formation show inhibition of Sterol 14-reductase, EBP, and CYP51

We next evaluated whether small-molecule promoters of oligodendrocyte formation identified using OPCs bearing leukodystrophy-causing mutations also inhibit cholesterol pathway enzymes. Pelizaeus-Merzbacher disease (PMD) is characterized by dysfunctional differentiation of OPCs to oligodendrocytes and subsequent hypomyelination; the “*jimpy*” (*Plp1*^jp^) mouse is frequently used to model PMD and shows a strong demyelinating phenotype.^23, 24^ Using iPSC-derived *jimpy* OPCs, a high-throughput screen identified a number of hits which were effective in promoting mutant oligodendrocyte formation.^11^ Notably, among the top 19 validated hit molecules identified using *jimpy* OPCs, eight have previously been shown to inhibit cholesterol pathway enzymes, including in wild-type mouse OPCs (Figure 4A).^15, 25^ Additionally, we established above that both Ro 25-6981 and fenspiride, which enhance oligodendrocyte formation in *jimpy* mice, are inhibitors of sterol 14-reductase and EBP, respectively (Figure 2C). We next evaluated the nine molecules not already linked to cholesterol pathway inhibition using our wild-type mouse OPC differentiation assay. We initially validated that five of these nine hit molecules promoted at least a 1.5-fold increase in MBP+ oligodendrocytes at the reported screening dose of 10 µM (Figure 4B and S3). The five oligodendrocyte-enhancing small molecules were then evaluated using GC-MS based sterol profiling to measure cholesterol pathway enzyme inhibition (Figure 4C). All five validated hits induced sterol accumulation at their effective screening dose, with three inhibiting EBP, one inhibiting sterol 14-reductase, and one inhibiting CYP51. Two of these molecules, climbazole (CYP51) and U-101958 (EBP), were also characterized at dose and showed comparable potency for enhancing both MBP+ oligodendrocyte formation and 8,9-unsaturated sterol levels (Figure 4D-G). Collectively, these data indicate that even small molecule enhancers of oligodendrocyte formation identified using OPCs harboring a disease-causing mutation are overwhelmingly likely to inhibit EBP, sterol 14-reductase, and CYP51.

**Figure 4.**
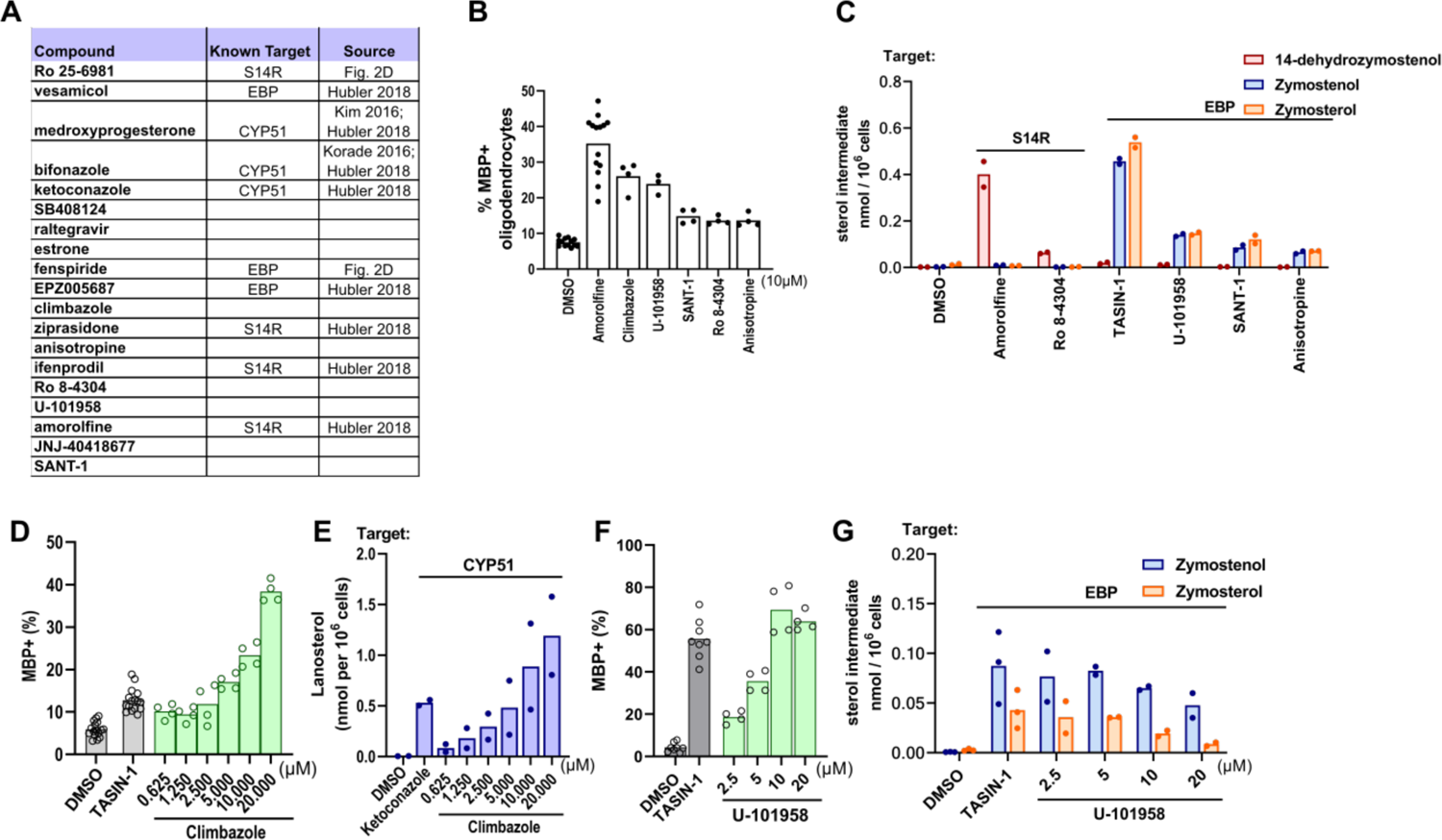
Many enhancers of oligodendrocyte formation inhibit cholesterol pathway enzymes. (A) Validated screening hits from Elitt et al., 2018 with annotations of subsequently-established inhibitory activity for EBP, CYP51, or sterol 14-reductase. (B) Percentage of MBP+ oligodendrocytes generated from mouse OPCs following treatment with bioactive small molecules at a uniform dose of 10 µM. *n*=4 wells per condition, except DMSO and amorolfine, *n*=16 wells, with >1,000 cells analyzed per well. (C) GC-MS based quantification of 14-dehydrozymostenol, zymostenol, and zymosterol levels in mouse OPCs treated for 24 h with the indicated screening hits at 10 µM. *n*=2 wells per condition. (D), (F) Percentage of MBP+ oligodendrocytes in mouse OPCs treated with climbazole and U-101958 in dose response, respectively. (E) GC-MS based quantification of lanosterol levels in mouse OPCs treated with climbazole in dose response as shown. (G) GC-MS based quantification of zymostenol and zymosterol levels in mouse OPCs treated with U-101958 in dose response as shown. *n*=2 wells per condition. For percentage of MBP+ oligodendrocytes for all bioactive small molecules that were screened, see Fig. S3. S14R, sterol 14-reductase.

### 2.4 Treatments effective in enhancing rat OPC differentiation inhibit EBP and sterol 14-reductase

Some past screens of bioactive small molecule libraries have used rat primary OPCs to identify enhancers of oligodendrocyte formation. We next tested whether leading enhancers of rat OPC differentiation identified by Deshmukh^14^ or Lariosa-Willingham^12^ promote formation of mouse oligodendrocytes and also inhibit EBP, sterol 14-reductase, or CYP51.

Deshmukh et al. identified 54 hits spanning 20 known mechanisms of action. Notably, five of these hits have subsequently been shown to inhibit EBP, sterol 14-reductase, or CYP51 in OPCs or other CNS cells (Figure 5A).^15, 25, 26^ Among the remaining hit molecules, we chose 10 with distinct known mechanisms of action for evaluation in our mouse OPC differentiation assay. Three molecules increased MBP+ oligodendrocytes at least 1.5-fold at the reported screening dose of 1.5 µM (Figure 5B); increasing the concentration to 4.5 µM did not result in additional hits (Figure S4B). Likewise, the screen reported by Lariosa-Willingham et al. identified 27 validated hits spanning seven drug classes. Remarkably, 11 of these hits, including all members of the ‘SERM’ and ‘Anti-fungal’ classes, have subsequently been shown to induce accumulation of 8,9-unsaturated sterols in OPCs or other CNS cells (Figure 5A).^15, 25^ We chose the most potent members of the remaining five drug class categories for evaluation in our mouse OPC differentiation assay; three of the five met our selection criteria for enhanced oligodendrocyte formation (Figure 5D).

**Figure 5.**
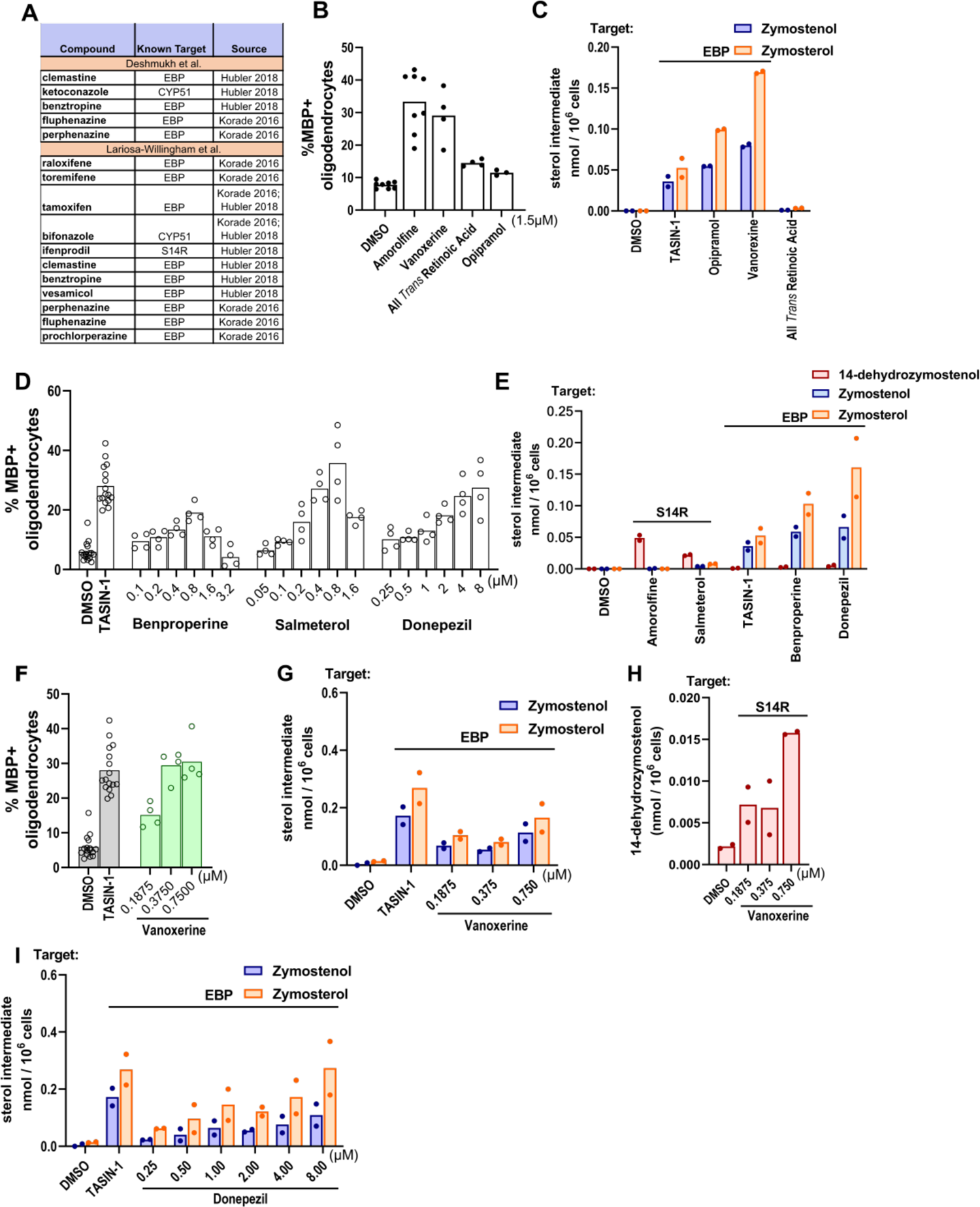
Top enhancers of both rat and mouse OPC differentiation inhibit EBP and sterol 14-reductase. (A) Validated screening hits from Deshmukh et al., 2013 and Lariosa-Willingham et al., 2016 with annotations of subsequently-established inhibitory activity for EBP, CYP51, or sterol 14-reductase. (B), (D) Percentage of MBP+ oligodendrocytes generated from mouse OPCs following treatment with bioactive small molecules. OPCs in (B) were treated with compounds at a uniform dose of 1.5 µM while those in (D) were treated in dose response as shown. *n=*4 wells per condition, except DMSO, amorolfine and TASIN-1, *n*=8 wells, with >1,000 cells analyzed per well. (C), (E) GC-MS based quantification of 14-dehydrozymostenol, zymostenol, and zymosterol levels in mouse OPCs treated for 24 h with the indicated screening hits at their effective concentrations. *n=*2 wells per condition. (F) Percentage of MBP+ oligodendrocytes in mouse OPCs treated with vanoxerine in dose response as shown. (G), (H), (I) GC-MS based quantification of zymostenol, zymosterol, and 14-dehydrozymostenol in mouse OPCs treated with vanoxerine and donepezil in dose response for 24 h. *n*=2 wells per condition. Results in (D) represent two independent experiments. For percentage of MBP+ oligodendrocytes for all bioactive small molecules that were screened, see Fig. S4. S14R, sterol 14-reductase.

The six validated hits identified in the Lariosa-Willingham and Deshmukh screens that promoted oligodendrocyte formation under our assay conditions, as well as three apparently inactive molecules (doxylamine, fasudil, and GW-1929), were then advanced to GC-MS based sterol profiling to determine whether these treatments modulate sterol levels in mouse OPCs (Figure 5C, 5E, 5G-I and S2C). Five of the six hits caused 8,9-unsaturated sterol accumulation at concentrations that enhanced oligodendrocyte formation, with four EBP inhibitors and one sterol 14-reductase inhibitor identified. All-*trans* retinoic acid (ATRA) was the only validated hit that did not cause sterol accumulation. Notably, donepezil and vanoxerine enhanced oligodendrocyte formation at concentrations that closely matched their potency for inducing accumulation of zymosterol and zymostenol (Figure 5F-I). Additionally, none of the three molecules tested that lacked effect in our OPC differentiation assay altered cellular sterol levels (Figure S2C). The identification of novel EBP or sterol 14-reductase inhibitory activity among molecules that robustly enhance differentiation of both rat and mouse OPCs further illustrates the high percentage of validated screening hits that induce accumulation of 8,9-unsaturated sterols to enhance oligodendrocyte formation.

### 2.5 Enhancers of human oligodendrocyte formation predominantly induce 8,9-unsaturated sterol accumulation

Recently the first high-throughput screen of human oligodendrocyte formation was reported using iPSC-derived human OPCs and a collection of 2500 bioactive small molecules.^27^ Remarkably, seven of 21 validated hits in this screen were previously demonstrated both to enhance formation of mouse oligodendrocytes and to induce accumulation of 8,9-unsaturated sterols in mouse OPCs.^15, 27^ While tamoxifen and imidazole antifungal drugs like oxiconazole have previously been shown to induce 8,9-unsaturated sterol accumulation in human cells, ^15, 21^ we confirmed that the remaining five hits also induced 8,9-unsaturated sterol accumulation in the human glioblastoma cell line GBM528 using our GCMS-based sterol profiling assay (Figure 6A-D). We then evaluated sterol levels after treatment with the remaining 14 hits from the screen not previously shown to modulate cholesterol biosynthesis and identified an additional three EBP inhibitors and two sterol 14-reductase inhibitors (Figure 6E-G). Collectively, 12 of 21 reported enhancers of human oligodendrocyte formation showed inhibition of EBP, CYP51, or sterol 14-reductase in a human CNS cell line. These 12 molecules also uniformly induced accumulation of 8,9-unsaturated sterols and promoted oligodendrocyte formation in mouse OPCs, further highlighting a high degree of correspondence between human and mouse OPC differentiation assays (Figure 6A-H and S5).^15, 21, 25, 27^ Together these findings support accumulation of 8,9-unsaturated sterols as a predominant phenotype among small molecules shown to enhance formation of oligodendrocytes from human OPCs.

**Figure 6.**
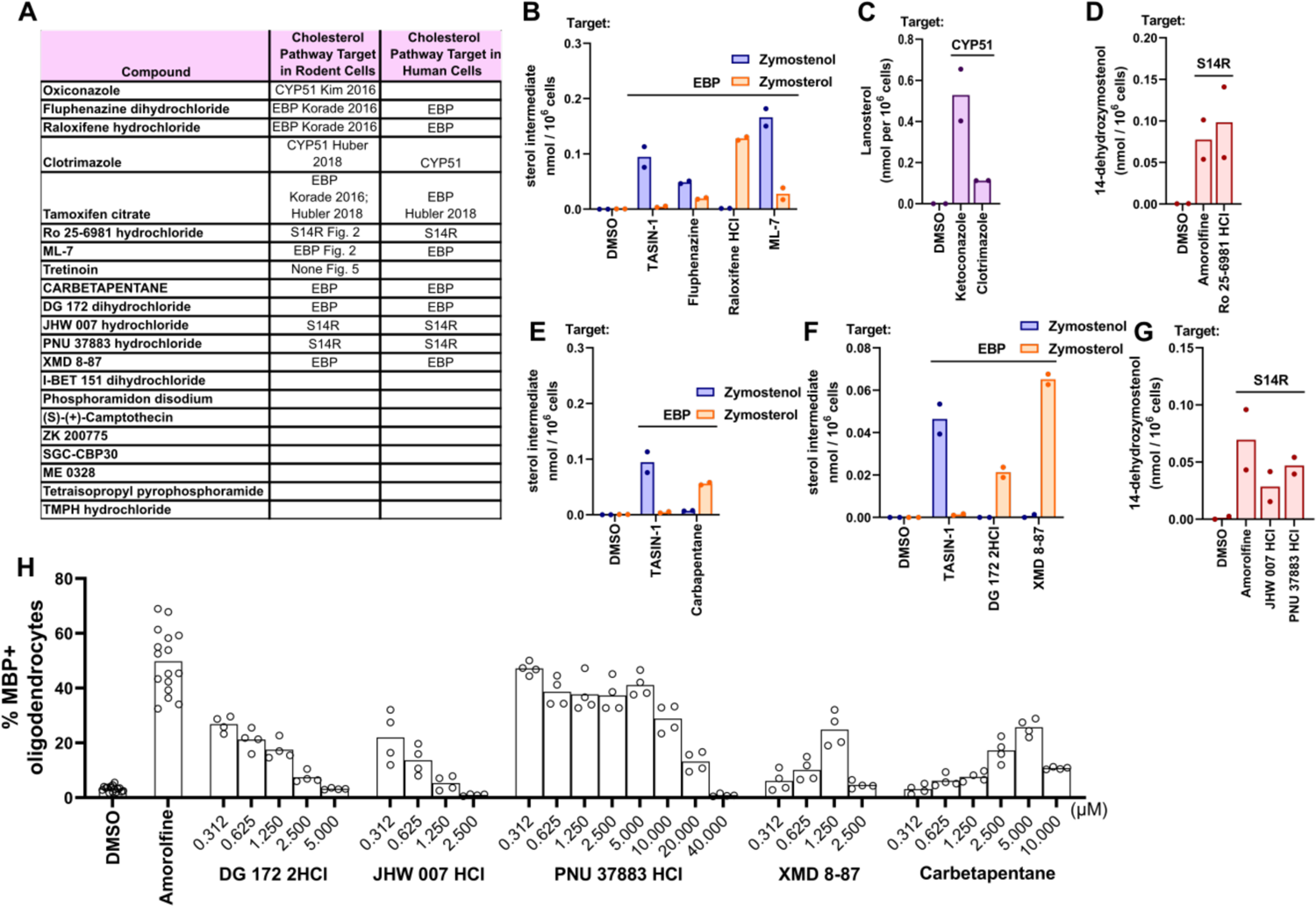
Top molecules reported to enhance human oligodendrocyte formation inhibit EBP, CYP51, and S14R in human GBM528 cells. (A) Validated screening hits from Li et al. 2022 with annotations of subsequently-established inhibitory activity for EBP, CYP51, or sterol 14-reductase. (B), (E), (F) GC-MS based quantification of zymostenol and zymosterol levels in GBM528 cells treated for 24 h with the indicated screening hits at 10 µM. (C) GC-MS based quantification of lanosterol levels in GBM528 cells treated for 24 h with clotrimazole at 10µM. (D), (G) GC-MS based quantification of 14-dehydrozymostenol levels in GBM528 cells treated for 24 h with the indicated molecules at 10 µM. *n*=2 wells per condition. (H) Percentage of MBP+ oligodendrocytes generated from mouse OPCs following treatment with bioactive small molecules in dose response as shown. *n=*4 wells per condition, except DMSO and amorolfine, *n*=8 wells with >1,000 cells analyzed per well.

### 2.6 Enhancers of oligodendrocyte formation advanced to clinical trials inhibit EBP

Several small-molecule enhancers of oligodendrocyte formation identified by high-throughput screening or other approaches have advanced into clinical investigation as potential multiple sclerosis therapies.^17, 28^ Two such molecules, clemastine and quetiapine, were first identified in a screen using rat oligodendrocyte wrapping of ‘micropillars’ as a primary readout.^13^ Previously, we confirmed the effects of clemastine as an enhancer of oligodendrocyte formation and established that clemastine inhibits EBP in mouse OPCs.^15^ Notably, quetiapine also enhanced oligodendrocyte formation and inhibited EBP in our assays (Figure 7A and 7D). Subsequently, bazedoxifene, a selective estrogen receptor modulator, was identified using the same micropillar-based screening platform.^29^ These studies discounted the estrogen receptor as bazedoxifene’s functional target in OPCs and used computational methods to suggest EBP as an alternative target. Indeed, GCMS-based sterol profiling revealed striking accumulation of the EBP substrate zymosterol in a dose-responsive manner parallel to bazedoxifene’s effect on oligodendrocyte formation, confirming bazedoxifene as an EBP inhibitor in mouse OPCs (Figure 7B and 7E). We next investigated GSK239512, a histamine receptor antagonist advanced clinically as a potential remyelinating therapeutic.^30^ We confirmed that GSK239512 enhanced oligodendrocyte formation at concentrations above 1 µM (Figure 7C). As other tertiary amine-containing histamine receptor modulators are known to inhibit EBP, we performed GCMS-based sterol profiling on mouse OPCs treated with GSK239512. Accumulation of EBP’s substrate sterols zymostenol and zymosterol indicated inhibition of EBP, and potency for EBP inhibition mirrored potency for enhancing oligodendrocyte formation (Figure 7F).

**Figure 7.**
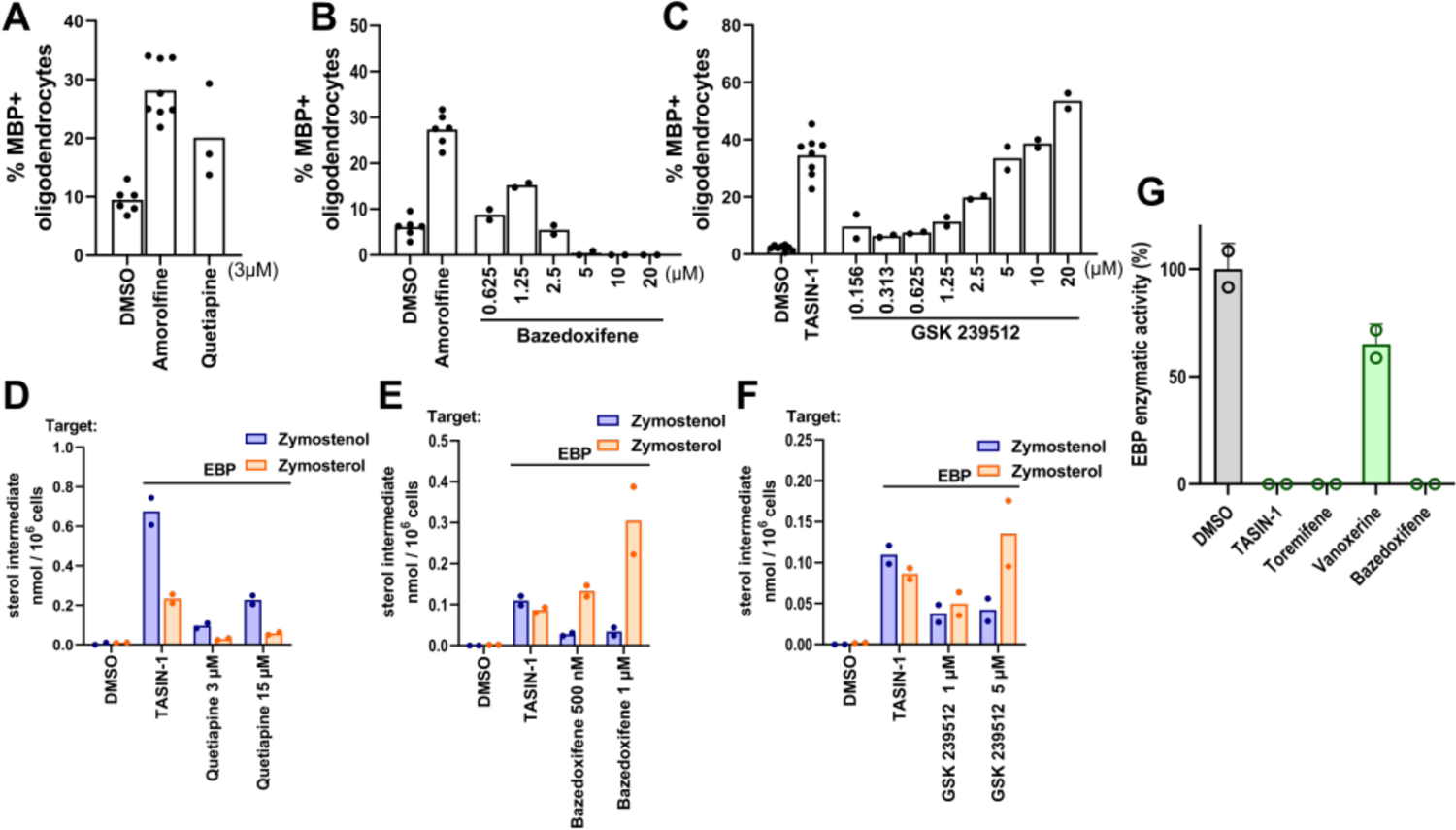
Compounds in clinical trials inhibit EBP. (A-C) Percentage of MBP+ oligodendrocytes generated from mouse OPCs following treatment with bioactive small molecules. Mouse OPCs in (A) were treated with quetiapine at 3 µM while those in (B) and (C) were treated with bazedoxifene and GSK 239512 in dose response as shown. *n=*4 wells per condition, except DMSO, amorolfine, and TASIN-1, *n*=8 wells, with >1,000 cells analyzed per well. (D-F) GC-MS based quantification of zymostenol, and zymosterol levels in mouse OPCs treated for 24 h with the indicated screening hits at their effective concentrations. *n=*2 wells per condition. (G) Quantification of EBP enzymatic activity in a biochemical assay. All treatments 10 µM. *n*=2 independent enzymatic assays. Bars indicate mean and error bars represent s.d. Results in (C) represent two independent experiments.

Finally, given the striking prevalence of EBP inhibitors found throughout this work, we performed a biochemical assay of EBP enzymatic activity using four top-performing molecules to evaluate whether these molecules directly inhibit EBP. While TASIN-1, toremifene, and bazedoxifene showed complete inhibition of EBP activity, vanoxerine showed partial inhibition; this result matches vanoxerine’s effects in cells where partial inhibition of both sterol 14-reductase and EBP is observed (Figure 7G, 5G-H). Altogether, these findings support EBP as the functional target by which bazedoxifene and other clinical remyelination candidates promote oligodendrocyte formation from cultured OPCs.

Our data highlight that an overwhelming fraction of enhancers of oligodendrocyte formation identified via HTS, and indeed several percent of molecules within these screening collections, target cholesterol biosynthesis enzymes that metabolize 8,9-unsaturated sterols. Recent studies by others have reinforced the high frequency with which FDA-approved drugs inhibit cholesterol pathway enzymes, including at doses used clinically.^26, 28, 31–34^ Available evidence suggests several explanations for why so many small molecules within available screening collections target cholesterol biosynthesis enzymes. First, many pathway enzymes metabolize cholesterol precursor sterols, which are highly hydrophobic substrates that occupy large and hydrophobic pockets within the active sites of these enzymes. Large hydrophobic binding pockets typically predict high enzyme ‘druggability’ with existing small molecule collections. Indeed, druggability analysis of recent X-Ray crystal structures of EBP using DoG SiteScorer revealed large binding pockets, a high fraction of hydrophobic residues within the pockets, and a subpocket druggability score as high as 0.84, well above the threshold of 0.5 typically used to predict a high level of druggability (Figure 8A).^33^ Similar findings were observed for CYP51, the only other enzyme that metabolizes 8,9-unsaturated sterols for which X-Ray crystal structure is available.^35^ As expected given the hydrophobic substrate binding pockets of pathway enzymes, hits validated in both our 2018 screen and our reanalysis in Figure 2 show significantly higher calculated log P than our screening collection as a whole (Figure 8B and 8C), indicating that more hydrophobic screening molecules are more likely to score as hits.^15^

**Figure 8.**
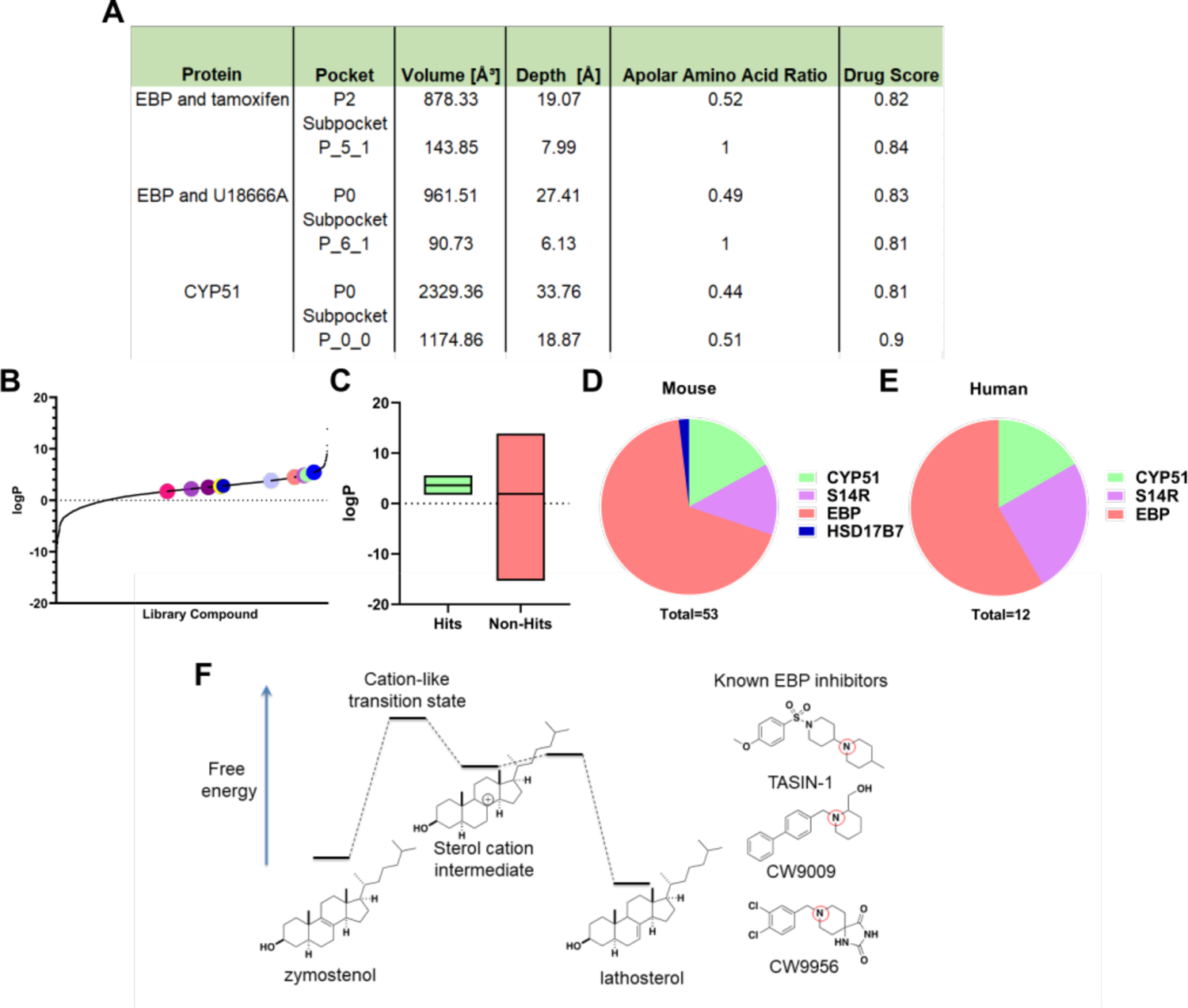
High logP hit molecules likely mimic sterol cation intermediate of highly druggable enzymes. (A) Drug Score analysis performed in DoGSiteScorer using crystal structures of EBP and tamoxifen, EBP and U18666A, and CYP51. (B) logP values for all library molecules tested. Dots represent the screening hits found in fig. 2 (A), (B). (C) Box plot comparing logP values for hit molecules and molecules that did not increase oligodendrocyte maturation. Included in the left box are our previously reported screening hits in addition to those found in fig. 2 (A), (B). (D) Total number of hit molecules that have been classified as either a CYP51, sterol 14-reductase, EBP, or HSD17B7 inhibitor in mouse OPCs. (E) Total number of molecules characterized as either a CYP51, sterol 14-reductase, or EBP inhibitor in human GBM528 cells. (F) Energy diagram comparing the structure of the sterol cation intermediate within EBP’s enzymatic process to known EBP inhibitors. Red circles indicate tertiary amines likely to be cationic at physiological pH. S14R, sterol 14-reductase.

While the presence of highly hydrophobic substrate binding pockets may generally predict high druggability among sterol-metabolizing enzymes, molecules that enhance oligodendrocyte formation are remarkably unequally spread among each of the 6 enzymes whose substrates are 8,9-unsaturated sterols. Including all hits from our 3,000 bioactive small molecules screened at 2 µM as well as those validated here or reported previously,^36^ we have characterized thirty-six EBP inhibitors, seven sterol 14-reductase inhibitors, nine CYP51 inhibitors, and 1 HSD17B7 inhibitor in mouse OPCs. Additionally, a similar distribution of seven EBP inhibitors, three sterol 14-reductase inhibitors, and one CYP51 inhibitor were characterized in human OPC-like glioblastoma cells (Figure 8D-E).^15^ No molecules targeting the remaining 8,9-unsaturated sterol-metabolizing enzymes, SC4MOL or NSDHL, have been identified in this or other screens, though inhibitors of SC4MOL that enhance oligodendrocyte formation were identified independent of screening efforts.^36^

The most commonly targeted enzyme, EBP, and sterol 14-reductase share enzymatic mechanisms that proceed through similar high-energy sterol cation intermediates.^33^ Notably, molecules identified as inhibiting EBP and sterol 14 reductase from screening libraries uniformly feature a tertiary amine functionality likely to be cationic at physiological pH. Potentially, these cationic amphiphilic molecules serve as ready mimics of the rate-limiting transition states of these enzymes, which by Hammond’s postulate likely closely resemble the high-energy sterol cation intermediate (Figure 8F). Evidence for this hypothesis includes the lack of known inhibitors of EBP and sterol 14-reductase that do not include a basic amine and past SAR studies showing that rendering the basic nitrogen non-basic via introduction of a carbonyl fully abrogates EBP inhibitory activity.^18^ Additionally, the two other sterol-metabolizing enzymes whose enzymatic mechanisms proceed via sterol cations, DHCR7 and DHCR24, while not targets for enhancing oligodendrocyte formation, have been disproportionately identified as off-target effects of FDA-approved drugs.^34^ These analyses suggest that while sterol-metabolizing enzymes likely share druggable hydrophobic pockets, enzymes whose mechanisms proceed via sterol cation intermediates are most likely to be targeted by currently available screening collections, which are enriched with cationic amphiphilic molecules. Importantly, many molecules within the cationic amphiphile class are inactive in our OPC differentiation and sterol profiling assays, highlighting that only those cationic amphiphiles that inhibit EBP or sterol 14-reductase enhance oligodendrocyte formation (Figure S2A-C).^15^

### 2.7 DISCUSSION

Previous work from our lab has shown that many small molecule enhancers of oligodendrocyte formation function through inhibition of several enzymes in the cholesterol biosynthesis pathway—CYP51, sterol 14-reductase, EBP, and more recently, LSS, SC4MOL, and HSD17B7.^15, 18, 36, 37^ As a result, the 8,9-unsaturated sterol substrates accumulate in OPCs and effectively enhance OPC differentiation and remyelination both *in vitro* and *in vivo*.^15 15^Though we have previously shown cholesterol biosynthesis enzyme inhibition to be a shared mechanism for several screening hits that enhance oligodendrocyte formation, the current study highlights the broad dominance of this mechanism in small molecule-driven oligodendrocyte formation. Whether assays use pluripotent stem cell-derived or primary OPCs, wild-type or *Jimpy* OPCs, or mouse, rat, or human OPCs, an overwhelming majority of validated hits inhibit cholesterol pathway enzymes and induce 8,9-unsatuarted sterol accumulation at concentrations than enhance oligodendrocyte formation. While we were unable to validate every reported hit molecule we tested in our assay, possibly due to differences in OPC source/species, differentiation assay conditions, or the expected emergence of false positives from screens, overall congruence among these screens is high. Especially noteworthy is the high fraction of hits identified in the first screen of human oligodendrocyte formation previously identified in past screens using rodent OPCs. Specific SERMs, anti-muscarinics, antifungals, antipsychotics, and ion channel modulators frequently emerge from multiple published screens (Figure 5A), providing confidence in the overall robustness and reproducibility of the OPC differentiation assay.^15^

Our findings add to the substantial body of recent work establishing that many FDA-approved drugs have potent off-target effects on cholesterol biosynthesis pathway enzymes. Of note, clomifene was identified as a novel EBP inhibitor in this investigation with a potent ability to enhance oligodendrocyte formation in the nanomolar range through EBP inhibition. As an orally available FDA-approved drug known to penetrate the blood-brain barrier,^38^ clomifene may be an attractive candidate for drug repurposing for treatment of demyelinating diseases.

Our interpretation of the dominance of cholesterol pathway modulators among known enhancers of oligodendrocyte formation is that a small percentage of cationic amphiphilic molecules, which are over-represented in available screening collections, can act as transition state mimics for EBP, sterol 14-reductase, and other sterol cation-metabolizing enzymes. This function is particularly likely in the low micromolar concentration range at which many high-throughput screens are run. Undoubtedly other targets and signaling pathways can also be modulated to drive oligodendrocyte formation; in particular, thyroid hormone receptor signaling is a well-validated axis confirmed in our present screen and known to be additive with sterol pathway modulators.^15^ Additionally, past and present work has noted enhanced oligodendrocyte formation for glucocorticoids and ATRA. Notably, in contrast to thyroid hormone, which promotes oligodendrocyte formation even at picomolar concentrations, glucocorticoids and ATRA affect oligodendrocyte formation only at concentrations near or above 1 µM. As these concentrations are far higher than those typically required to modulate the glucocorticoid receptor or retinoic acid receptor, the cellular targets mediating the effects of these molecules on OPCs are uncertain. Doubtless additional druggable targets beyond these remain to be established for enhancing oligodendrocyte formation, and future small-molecule screens that take steps to rapidly triage or suppress the large number of cholesterol pathway inhibitors likely to be present in typical screening collections may be more likely to identify such novel targets.

## 3. MATERIALS AND METHODS

### 3.1 OPC Media Conditions

*OPC growth media* refers to media containing DMEM/F-12 media (ThermoFisher, 11320-033) supplemented with 1X N2Max, 1X B27 (ThermoFisher), FGF2 (10 µg/mL, R&D systems, 233-FB025), and PDGF-AA (10 µg/mL, R&D systems, 233-AA-050). *Differentiation-permissive media* refers to the same media conditions as OPC growth media with the exclusion of FGF and PDGF.

### 3.2 Mouse OPC Preparation

iPSC-derived OPCs were gifted from the Tesar Lab and were previously obtained using in-vitro differentiation protocols and cell culture conditions (Case Western Reserve University).^39^ OPCs were expanded and subsequently cryopreserved at −80°C using gradually decreasing temperatures (1°C/min). Cryopreserved OPCs were stored in liquid nitrogen before further handling. OPCs were thawed into OPC growth media with 1X Penicillin/Streptomycin (Sigma) for one passage before the use of the OPCs in assays. The OPC cell cultures were regularly tested to be mycoplasma free using a MycoAlert Mycoplasma Detention Kit (Lonza, LT07-118). The OPCs were authenticated with immunopositivity for OPC cell markers (Nkx2—2, Olig2, and Sox10). Finally, OPCs were identified for their ability to differentiate into mature, MBP positive oligodendrocytes in differentiation-permissive media in the presence of thyroid hormone.

### 3.3 Cell Culture

Before seeding OPCs for assays, OPCs were expanded in 25 mL of OPC growth media for 4 days, with replenishment of new media every 2 days, in OPC growth media in T175 flasks coated with poly-ornithine (Sigma) and laminin (Sigma). Coating was performed by incubating 25 mL of a 15 µg/mL poly-ornithine solution in water at 37°C for 1 h before completely drying the flask. Then, the flasks were coated with 12 mL of a 15 µg/mL laminin in PBS solution at 37°C for 1 h before aspirating and the addition of cells and OPC growth media to each flask. The human glioma cell line GBM528 was a gift of Jeremy Rich (Cleveland Clinic). GBMs were expanded in T175 flasks in DMEM/F-12 media (ThermoFisher, 11320-033) supplemented with N2 Max, B27 (ThermoFisher), 20 ng/ml FGF and 20 ng/ml EGF. Cells were non-adherent so passaging and plating cells involved initially spinning down cells and subsequently incubating cells at 37℃ with accutase to break up apparent colonies of cells.

### 3.4 OPC Phenotypic Assay for MBP Positivity

Following initial expansion of OPCs (see OPC Cell Culture), OPCs were seeded onto 96-well CellCarrierUltra plates (PerkinElmer) coated with poly-D-lysine (Sigma; 15 µg/mL) and laminin (Sigma; 15 µg/mL) at 37°C for 1 h each. For each 96-well plate, 40,000 cells were seeded in each well. Cells were incubated at 37°C for 72 h and subsequently fixed with 4% paraformaldehyde in PBS for 20 min. The fixed plates were washed with PBS (200 µL per well) three times and then permeabilized and blocked using 0.1% Triton X-100 (Sigma) with 10% donkey serum (v/v) in PBS for 1 h. Cells were labelled with an anti-MBP antibody (ABCAM, ab7349; 1:200) for 16 h at 4°C followed by secondary antibody staining with Alexa Fluor 488 donkey-anti rat (1:500) for 45 min. DAPI staining was added for nuclei visualization (Sigma; 1 µg/mL). After both primary and secondary antibody staining, cells were washed three times with PBS (200 µL) using a multi-channel pipette and a 96-well aspiration manifold (Biotek).

### 3.5 100 nM Bioactive Small Molecule Library Screen

Following expansion of OPCs (see OPC Cell Culture), OPCs were seeded onto 384-well CellCarrierUltra plates (PerkinElmer) coated with poly-D-lysine (15 µg/mL; 37°C for 1 h) and laminin (15 µg/mL; 37°C for 1 h) at a concentration of 10,000 cells per well in 50 µL of differentiation-permissive media. An EL406 Microplate Washer Dispenser (Biotek) equipped with a 5 µL dispensing cassette (Biotek) was used to dispense cells. A 100 µM stock of each drug in the bioactive library (Selleck Bioactive Compound Library-I; L1700) dissolved in dimethyl sulfoxide (DMSO) was prepared in Abgene storage 384-well plates (ThermoFisher Scientific). Drugs were added to the plates using a 50 nL solid pin tool attached to a Janus automated workstation (PerkinElmer), for a final screening concentration of 100 nM. DMSO was run as a negative control for each plate. A previously reported OPC differentiation-enhancing drug, TASIN-1, was used as a positive control at a concentration of 1 µM.^15^ Cells were incubated at 37°C for 72 h and subsequently fixed and stained for MBP and DAPI according to the protocol in OPC Phenotypic Assay for MBP Positivity. Imaging and analysis of the plates was performed as outlined in High-Content Imaging and Analysis.

### 3.6 High-Content Imaging and Analysis

Following fixing and staining OPC phenotypic assays for MBP positivity, plates were imaged on the Operetta High-Content Imaging and Analysis system (PerkinElmer). A set of 6 fields were captured in each well of each 96-well plate (4 fields for 384-well plates) resulting in an average of 1600 cells (500 cells for 384-well plate) scored per well. Analysis was performed using PerkinElmer Harmony and Columbus software. First, live nuclei were identified using the DAPI channel with a traced nucleus area between 50-250 µm^2^ which excludes pyknotic nuclei.^11, 15, 18^ The traced nucleus region was then expanded by 50% and examined for myelin basic protein (MBP) staining to identify mature oligodendrocyte nuclei.^15^ Dividing the number of mature oligodendrocyte nuclei by the number of live cells identified yielded % MBP positive oligodendrocytes.

### 3.7 Gas Chromatography–Mass Spectrometry Sterol Profiling

OPCs were plated at a density of 1M cells per well (2 mL) in poly-ornithine (15 µg/mL; 37°C for 1 h) and laminin (15 µg/mL; 37°C for 1 h) coated six-well plates (Costar) with differentiation-permissive media. After 24 h, cells were rinsed with PBS in the plate and then the plate was frozen at −80°C. For analysis of sterols, cells were lysed using 1 mL of hexane (Sigma-Aldrich) for 10 min. 100 µL of cholesterol-d7 standard in hexane (Cambridge Isotope Laboratories) was added to each biological sample before drying under nitrogen. Following this, 55 µL of bis(trimethylsilyl)trifluoroacetamide/ trimethylchlorosilane was added to form trimethylsilyl sterol derivatives by heating at 60°C for 20 min. Following this sterol derivatization, 1 µL was analyzed by gas chromatography-mass spectrometry using an Agilent 7890B GC system. The following *m/z* ion fragments were used to quantitate each metabolite: cholesterol-d7 (465), cholesterol (368), zymosterol (456), zymostenol (458), desmosterol (343), lanosterol (393), lathosterol (458), FF-mas (482), and 14-dehydrozymostenol (456). Standard curves were generated by injecting concentrations of sterol standards with a fixed amount of cholesterol-d7.^15^ This protocol was also used for GC-MS based sterol profiling of GBM528 cells.

### 3.8 EBP enzymatic assay

EBP enzymatic activity was measured using a reported method with slight modifications^22^: mouse microsomes containing active EBP was obtained, small molecule treatments were added at their corresponding concentrations, and zymostenol was added to a final concentration of 25 µM in a final reaction volume of 500 μL. The reaction was then incubated at 37°C for 2 h. Sterols were extracted using hexane, cholesterol-d7 was added to enable quantitation, and the pooled organics were dried and evaporated under nitrogen gas. Samples were then silylated and analyzed using GC/MS (see Gas Chromatography–Mass Spectrometry Sterol Profiling).

### 3.9 Statistical Analysis

No statistical methods were used to predetermine sample size. Number of replicates and independent experiments performed are noted in the figure legend. P-values were calculated with tests indicated in the figure legends. Graphpad Prism 9.2.0 was used to perform all statistical analysis.

## Supporting information

Five Supporting Figures

## AUTHOR INFORMATION

### Corresponding Author

**Drew J Adams --** *Department of Genetics and Genome Sciences, Case Western Reserve University School of Medicine, Cleveland, Ohio 44106, USA*.

## Authors

**Joel L Sax --** *Department of Genetics and Genome Sciences, Case Western Reserve University School of Medicine, Cleveland, Ohio 44106, USA*.

**Samantha N Hershman --** *Department of Genetics and Genome Sciences, Case Western Reserve University School of Medicine, Cleveland, Ohio 44106, USA*.

**Zita Hubler --** *Department of Genetics and Genome Sciences, Case Western Reserve University School of Medicine, Cleveland, Ohio 44106, USA*.

**Dharmaraja Allimuthu --** *Department of Genetics and Genome Sciences, Case Western Reserve University School of Medicine, Cleveland, Ohio 44106, USA*.

**Matthew S Elitt --** *Department of Genetics and Genome Sciences, Case Western Reserve University School of Medicine, Cleveland, Ohio 44106, USA*.

**Ilya Bederman --** *Department of Genetics and Genome Sciences, Case Western Reserve University School of Medicine, Cleveland, Ohio 44106, USA*.

## Author Contributions

J.S., S.H., Z.H, D.A., M.E., and D.J.A. evaluated the effects of small molecules on *in vitro* oligodendrocyte formation. J.S., S.H., Z.H., D.A., M.E., I.B., and D.J.A. performed and analyzed the OPC gas chromatography-mass spectrometry sterol profiling experiments *in vitro*. S.H. and D.J.A. analyzed druggability of EBP and CYP51 and lipophilicity trends of molecules within the screening collection. J.S., S.H., and D.J.A. wrote the manuscript. All authors provided intellectual input, edited, and approved the final manuscript.

## Notes

The authors declare the following competing interests: D.J.A. is a founder, consultant, director, and shareholder of Convelo Therapeutics, Inc., which seeks to develop remyelinating therapeutics. D.J.A is also an inventor on patents and patent applications that have been licensed to Convelo Therapeutics, Inc.

## Supporting Information

The Supporting Information is available free of charge at http://pubs.acs.org. The Supporting Information containing Figures S1 to S5 include additional OPC differentiation and GC-MS-based sterol profiling results to support the study.

## Acknowledgements

The authors thank P. Tesar for providing OPCs and reagents. Y. Fedorov, the CWRU Small-Molecule Drug Development Core, and M. Drumm are thanked for technical support. Table of contents graphic generated using Biorender.com.

---

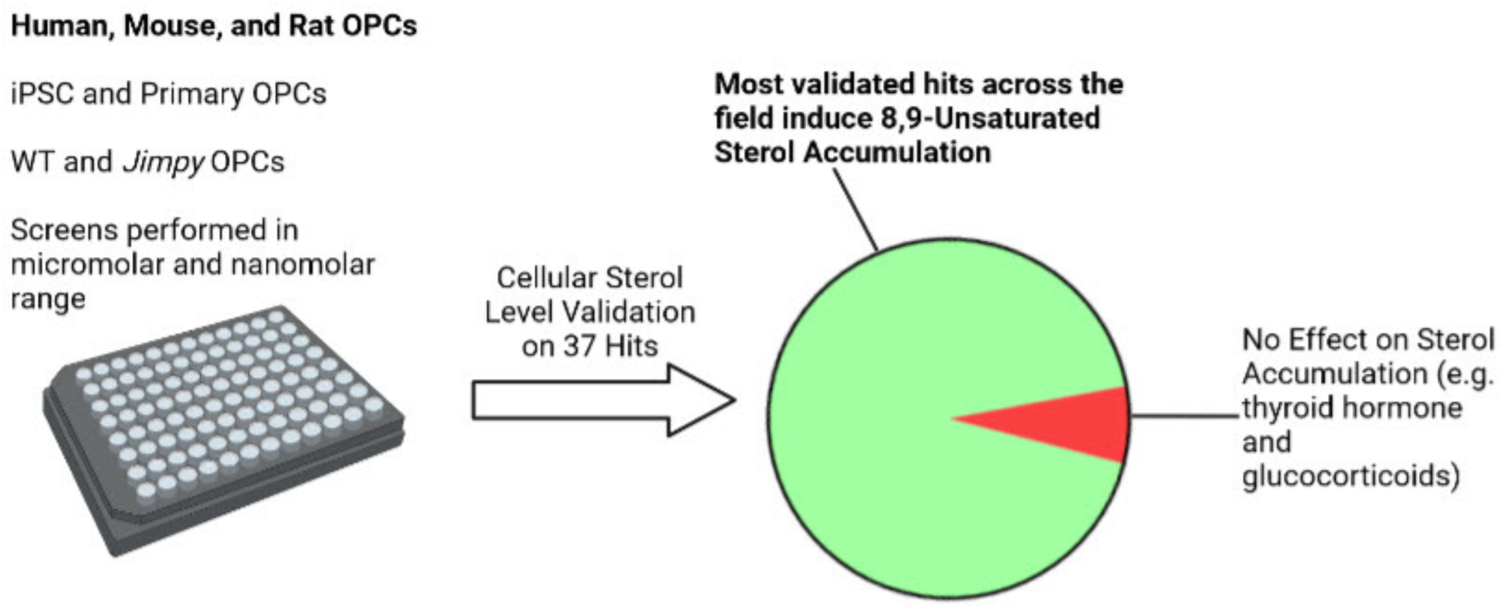

