## Supplementary material for "Enhancers of human and rodent oligodendrocyte formation predominantly induce cholesterol precursor accumulation": Five Supporting Figures

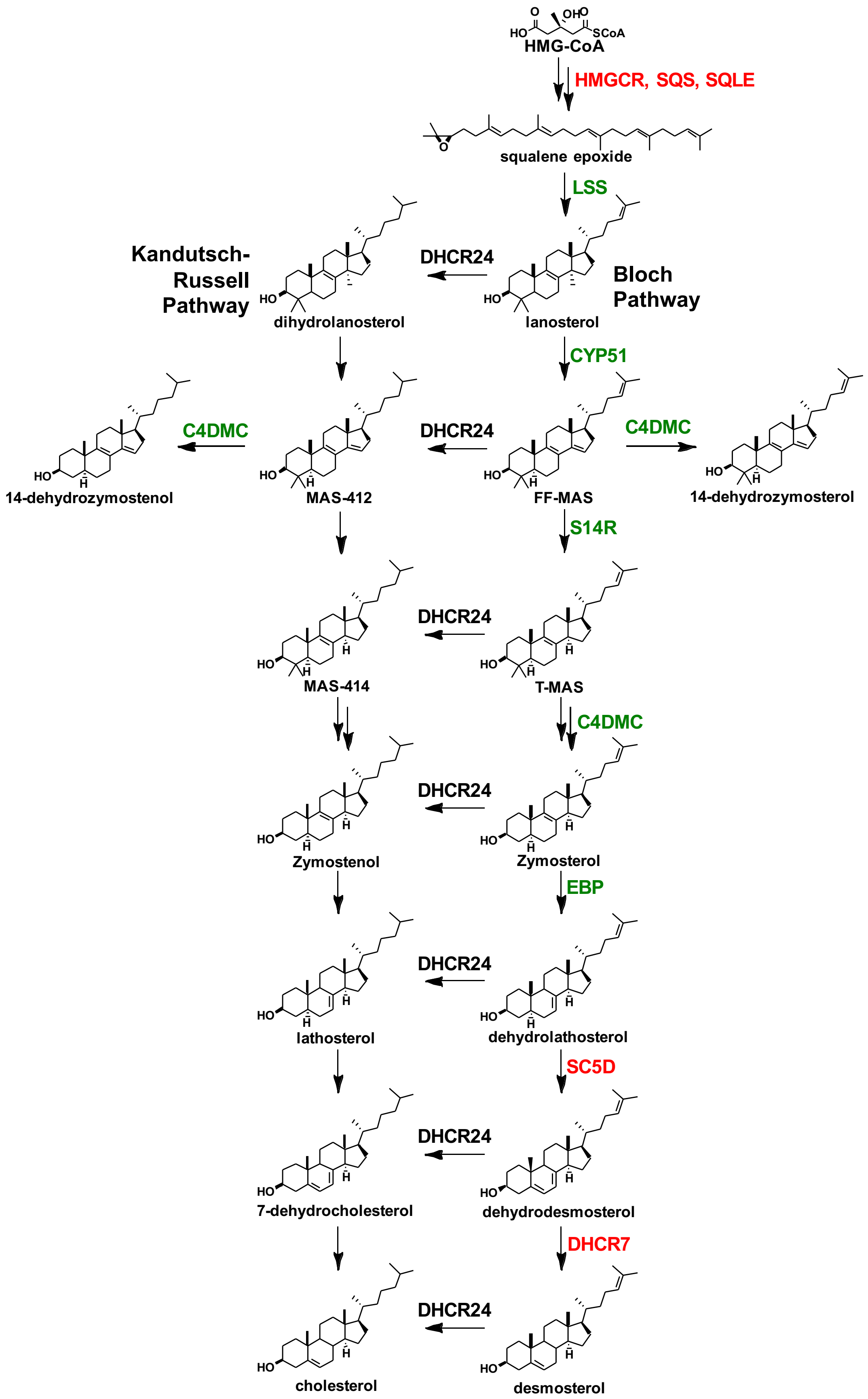
HMG-CoA
HMGCR, SQS, SQLE
squalene epoxide
LSS
Kandutsch-Russell Pathway
Bloch Pathway
dihydrolanosterol
DHCR24
lanosterol
CYP51
14-dehydrozymostenol
C4DMC
MAS-412
DHCR24
FF-MAS
C4DMC
14-dehydrozymosterol
S14R
MAS-414
DHCR24
T-MAS
C4DMC
Zymostenol
DHCR24
Zymosterol
EBP
lathosterol
DHCR24
dehydrolathosterol
SC5D
7-dehydrocholesterol
DHCR24
dehydrodesmosterol
DHCR7
cholesterol
DHCR24
desmosterol

**Figure S1| Cholesterol biosynthesis pathway diagram** Cholesterol biosynthesis pathway diagram highlighting the Kandutsch-Russel and Bloch pathways. Highlighted in green are enzymes in the pathway that when inhibited lead to enhanced oligodendrocyte formation. Enzymes highlighted in red are enzymes that do not increase oligodendrocyte formation when inhibited.

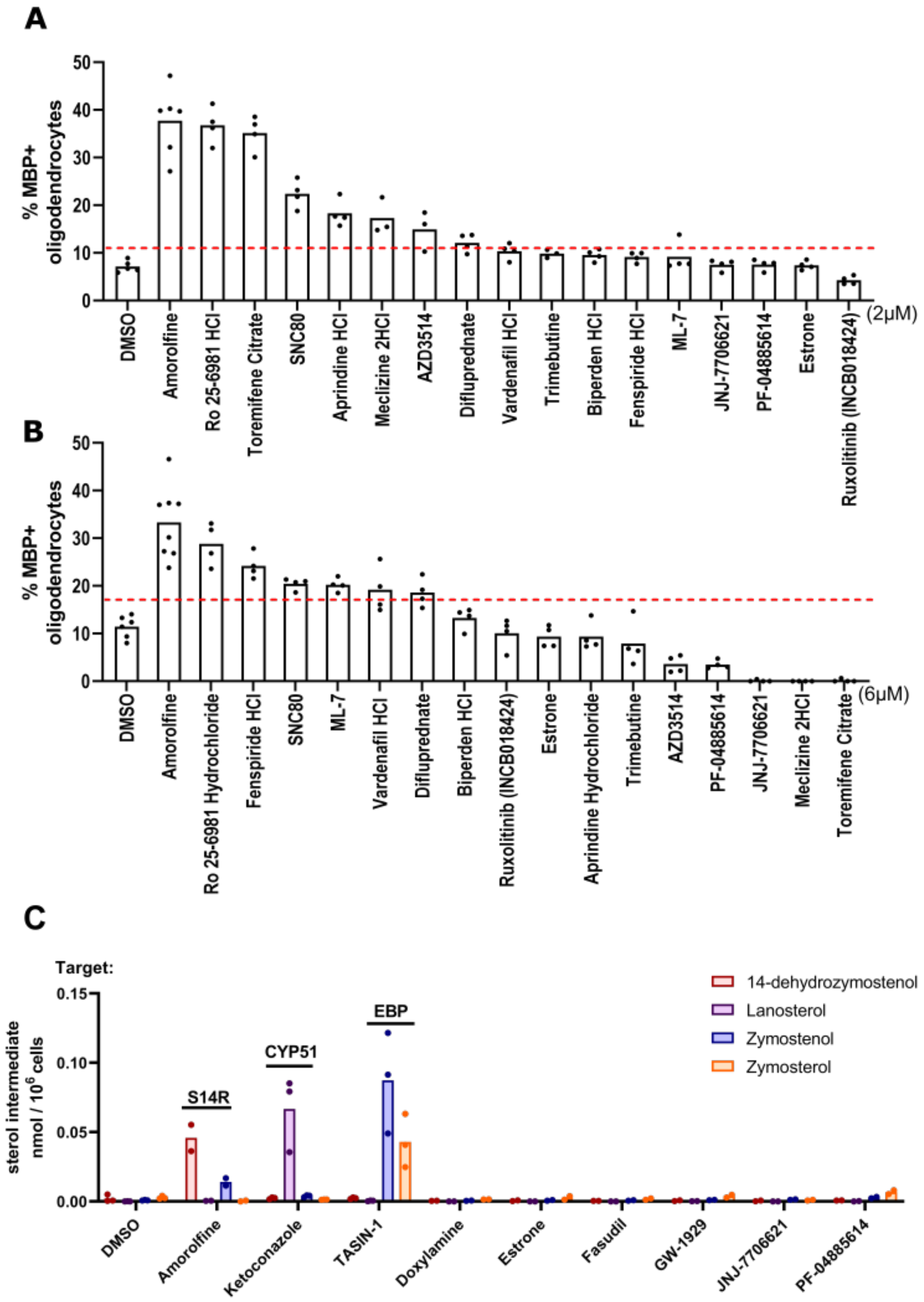
A
% MBP+ oligodendrocytes
0
10
20
30
40
50
DMSO
Amorolfine
Ro 25-6981 HCl
Toremifene Citrate
SNC80
Aprindine HCl
Meclizine 2HCl
AZD3514
Difluprednate
Vardenafil HCl
Trimebutine
Biperden HCl
Fenspiride HCl
ML-7
JNJ-7706621
PF-04885614
Estrone
Ruxolitinib (INCB018424)
(2μM)
B
% MBP+ oligodendrocytes
0
10
20
30
40
50
DMSO
Amorolfine
Ro 25-6981 Hydrochloride
Fenspiride HCl
SNC80
ML-7
Vardenafil HCl
Difluprednate
Biperden HCl
Ruxolitinib (INCB018424)
Estrone
Aprindine Hydrochloride
Trimebutine
AZD3514
PF-04885614
JNJ-7706621
Meclizine 2HCl
Toremifene Citrate
(6μM)
C
Target:
sterol intermediate nmol / 10^6 cells
0.00
0.05
0.10
0.15
S14R
CYP51
EBP
14-dehydrozymostenol
Lanosterol
Zymostenol
Zymosterol
DMSO
Amorolfine
Ketoconazole
TASIN-1
Doxylamine
Estrone
Fasudil
GW-1929
JNJ-7706621
PF-04885614

**Figure S2| a, b,** Percentage of MBP+ oligodendrocytes generated from mouse OPCs following treatment with bioactive small molecules at uniform doses of 2µM (**a**) and 6µM (**b**). *n*=4 wells per condition, except DMSO and amorolfine, a known S14R inhibitor, *n*=8 wells, with >1,000 cells analyzed per well. Dashed red lines represent the DMSO average of % MBP+ oligodendrocytes multiplied by 1.5 (**a**=10.70, **b**=17.16). **c,** GC-MS based quantification of 14-dehydrozymostenol, lanosterol, zymostenol, and zymosterol in mouse OPCs treated for 24 h with known cholesterol biosynthesis inhibitors and six compounds shown to be inactive in the differentiation assay. Cationic, amphiphilic compounds include fasudil, doxylamine, and PF-04885614. Non-cationic, amphiphilic molecules include JNJ-7706621, GW-1929, and estrone. S14R= sterol-14 reductase.

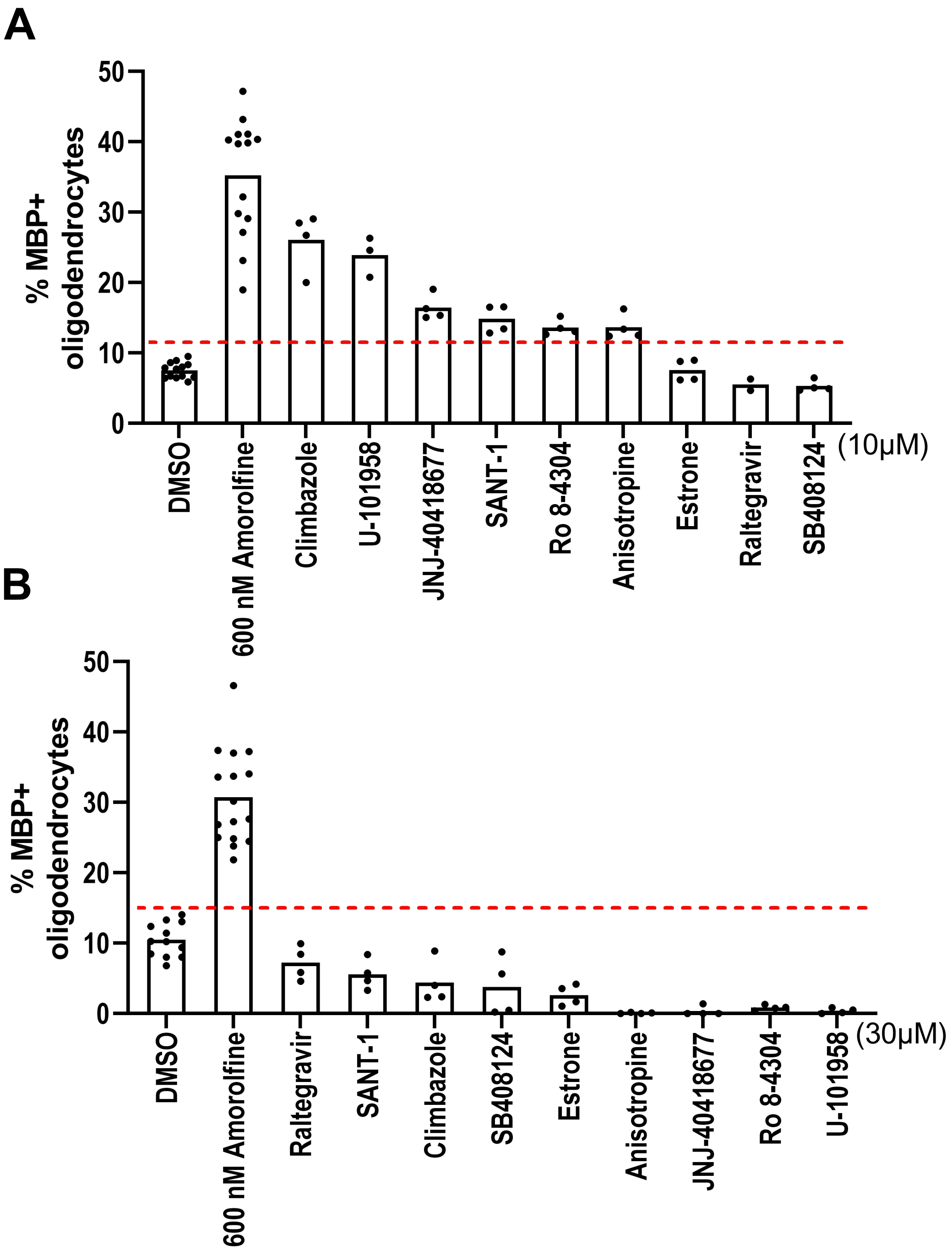


**Figure S3| a, b,** Percentage of MBP+ oligodendrocytes generated from mouse OPCs following treatment with bioactive small molecules at uniform doses of 10μM (**a**) and 30μM (**b**). *n*=4 wells per condition, except DMSO and amorolfine, a known S14R inhibitor, *n*=8 wells, with >1,000 cells analyzed per well. Dashed red lines represent the DMSO average of % MBP+ oligodendrocytes multiplied by 1.5 (**a**=11.29, **b**=15.69).

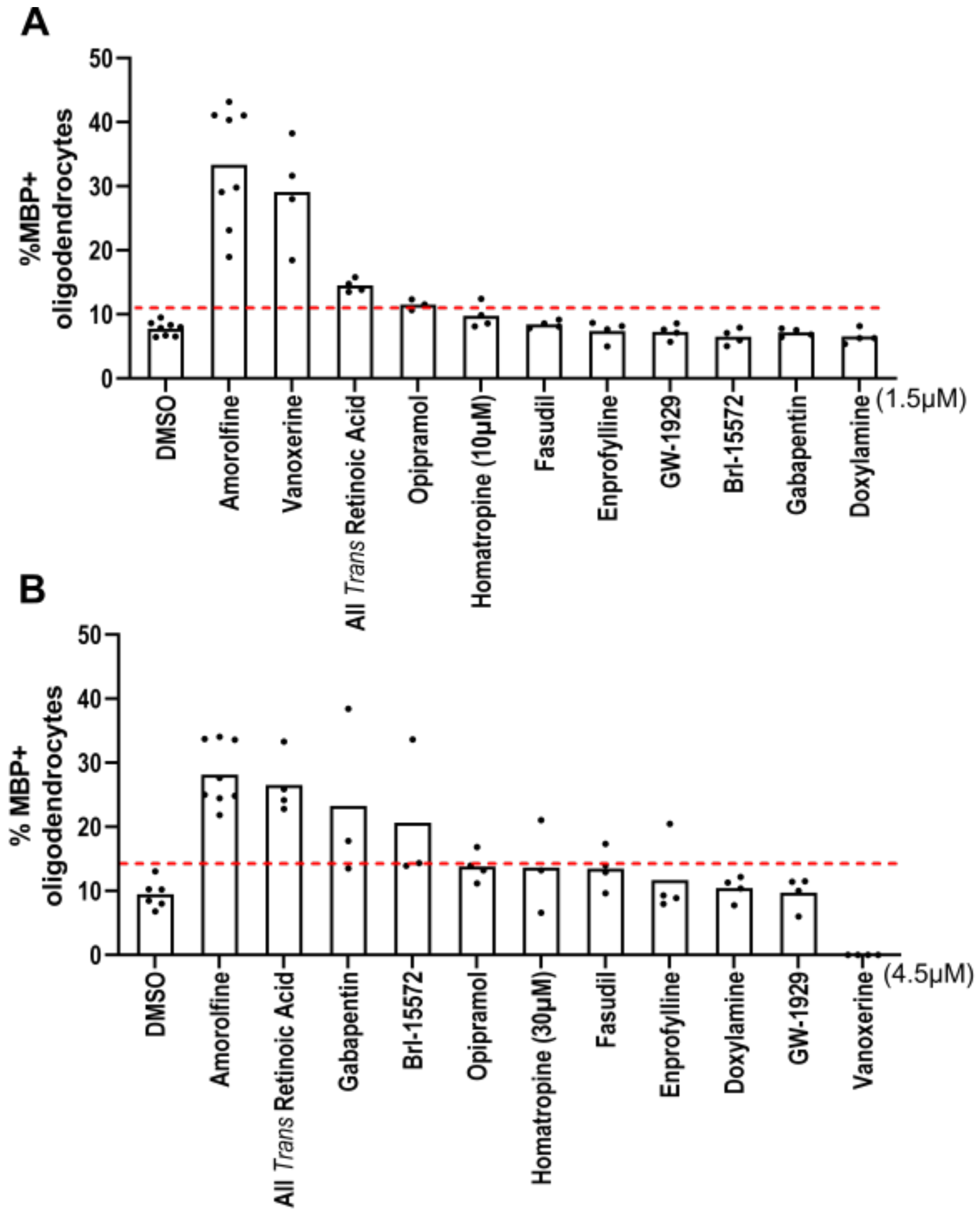


**Figure S4| a, b,** Percentage of MBP+ oligodendrocytes generated from mouse OPCs following treatment with bioactive small molecules at uniform doses of 1.5µM (**a**) and 4.5µM (**b**). *n*=4 wells per condition, except DMSO and amorolfine, a known S14R inhibitor, *n*=8 wells, with >1,000 cells analyzed per well. Dashed red lines represent the DMSO average of % MBP+ oligodendrocytes multiplied by 1.5. (**a**=11.66, **b**=14.22).

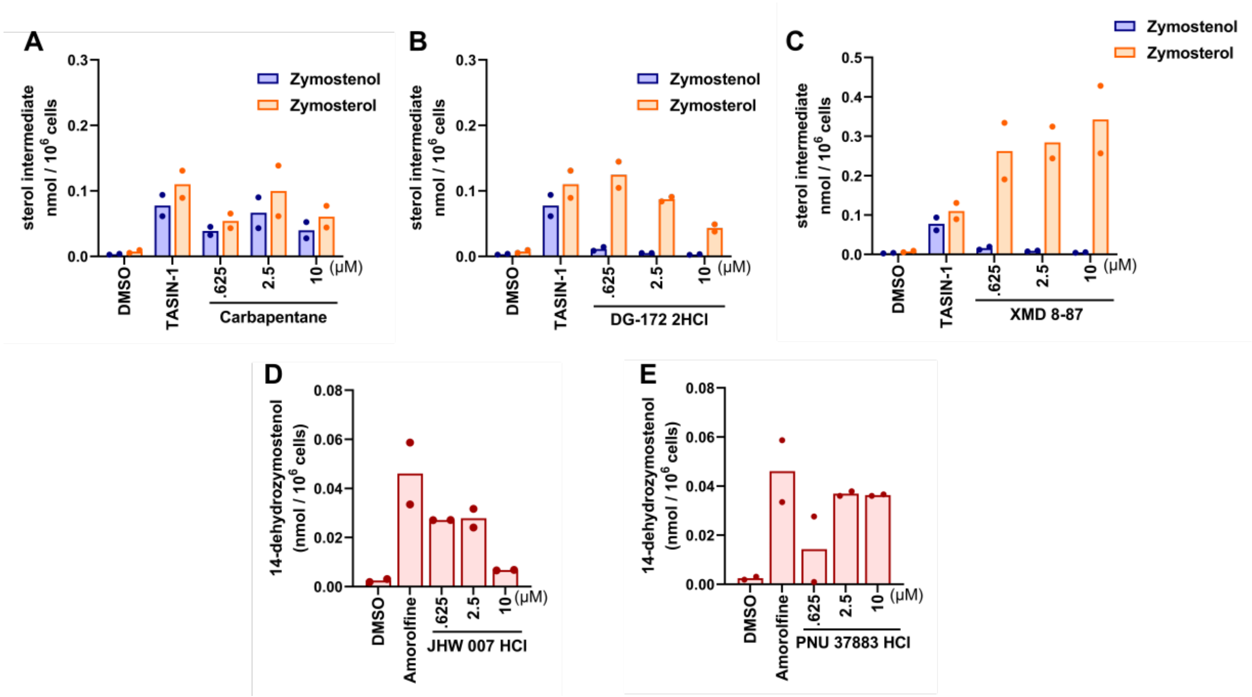


**Figure S5| a, b, c,** GC-MS based quantification of zymostenol and zymosterol levels in mouse OPCs treated for 24 h with the indicated screening hits in dose response as shown. *n*=2 wells per condition. **d, e,** GC-MS based quantification of 14-dehydrozymostenol levels in mouse OPCs treated for 24 h with the indicated screening hits in dose response as shown. *n*=2 wells per condition
